# Exploring the complexity of soybean (*Glycine max*) transcriptional regulation using global gene co-expression networks

**DOI:** 10.1101/2020.06.19.161950

**Authors:** Fabricio Almeida-Silva, Kanhu C. Moharana, Fabricio B. Machado, Thiago M. Venancio

**Affiliations:** Laboratório de Química e Função de Proteínas e Peptídeos, Centro de Biociências e Biotecnologia, Universidade Estadual do Norte Fluminense Darcy Ribeiro, Campos dos Goytacazes, RJ, Brazil

**Author notes:** TMV: Laboratório de Química e Função de Proteínas e Peptídeos, Centro de Biociências e Biotecnologia, Universidade Estadual do Norte Fluminense Darcy Ribeiro. Av. Alberto Lamego 2000, P5, sala 217, Campos dos Goytacazes, RJ, Brazil.

**Keywords:** transcription factors, gene duplication, gene function, genome evolution, systems biology

## Abstract

Soybean (*Glycine max* (L.) Merr.) is one of the most important crops worldwide, constituting a major source of protein and edible oil. Gene co-expression networks (GCN) have been extensively used to study transcriptional regulation and evolution of genes and genomes. Here, we report a soybean GCN using 1,284 publicly available RNA-Seq samples from 15 distinct tissues. We found modules that are differentially regulated in specific tissues, comprising processes such as photosynthesis, gluconeogenesis, lignin metabolism, and response to biotic stress. We identified transcription factors among intramodular hubs, which probably integrate different pathways and shape the transcriptional landscape in different conditions. The top hubs for each module tend to encode proteins with critical roles, such as succinate dehydrogenase and RNA polymerase subunits. Importantly, gene essentiality was strongly correlated with degree centrality and essential hubs enriched in genes involved in nucleic acids metabolism and regulation of cell replication. By using a using a guilt-by-association approach, we predicted functions for 93 of 106 hubs without functional description in soybean. Most of the duplicated genes had different transcriptional profiles, supporting their functional divergence, although paralogs originating from whole-genome duplications (WGD) are more often preserved in the same module than those from other mechanisms. Together, our results highlight the importance of GCN analysis in unraveling key functional aspects of the soybean genome, in particular those associated with hub genes and WGD events.

## INTRODUCTION

Soybean (*Glycine max* (L.) Merr.) is one of the most important legume crops worldwide, being important for human nutrition, animal feed, and biotechnological applications. It has a paleopolyploid genome that resulted from two whole-genome duplication (WGD) events that happened 58 and 13 million years ago [1]. Because of these WGD events, ∼75% of the soybean genes are present in multiple copies [2]. Over the last decade, several transcriptomic studies have investigated soybean transcriptional programs in different tissues and conditions [3-6]. However, most works explore only a few conditions and developmental stages, typically using pathway/function enrichment methods. Such approaches can overlook a myriad of interesting transcriptional patterns, which could otherwise be unraveled by integrative studies using hundreds of samples submitted to systems-level analyses.

A powerful approach to further explore transcriptomic data is the use of co-expression networks. Co-expression networks are undirected graphs comprising nodes representing genes that are connected by edges whenever significantly co-expressed. Co-expression networks have been used to understand transcriptional regulation in many plant species, such as maize (*Zea mays* L.), rice (*Oryza sativa* L.), tomato (*Solanum lycopersicum* L.), thale cress (*Arabidopsis thaliana* (L.) Heynh), and soybean [7-10]. However, most co-expression network analyses focus on case-control experiments, limiting their potential to uncover broader biological patterns [11-13]. A recent global co-expression network analysis of 1,072 soybean microarray samples revealed a gene module that is likely involved in the evolution of nodulation in plants [14]. In spite of these interesting results, hybridization-based transcriptional quantification often suffer from poor reproducibility [15]. Over the past few years, data from hundreds of soybean RNA-seq samples have accumulated and were recently integrated by our group in a large and publicly available soybean transcriptome atlas [16]. Nevertheless, a network analysis using this collection remains to be performed.

Co-expression analyses have recently been used to study the evolution of gene function in plants [17, 18]. After a duplication event, selective constraints are relaxed and gene copies typically undergo two possible fates: i. gene fractionation, when one copy accumulate loss-of-function mutations and eventually erode from the genome and; ii. gene retention via subfunctionalization or neofunctionalization [19]. Although pseudogenization is the most common fate of duplicated genes [20], genes encoding regulatory proteins (e.g. transcription factors – TFs – and signal transduction proteins) or subunits of protein complexes (e. g. proteasome and ribosome) are retained more often than expected [21, 22], mostly because of the maintenance of their stoichiometry [21]. However, in case of local duplications (e.g. tandem duplications), TFs and subunits of protein complexes tend to be fractionated because of dosage sensitivity [23].

Gene retention is thought to be more likely if the newly duplicated genes remain in the genome for an extended window of time, allowing mutation and selection pressures to rewire gene functions [24]. The vast majority of retained genes undergo subfunctionalization and, more rarely, neofunctionalization [25]. At the transcriptional level, subfunctionalization of a gene pair typically happens through mutations at cis-regulatory regions, resulting in two possible scenarios: i. both genes reduce their transcription, making them both necessary to perform the original function or; ii. genes from the pair become expressed in different tissues or conditions (transcriptional divergence) [26]. Thus, a network-level gene expression analysis could shed light on the possible fates of duplicated genes.

Here, we report a GCN built with data from 1,284 RNA-seq samples to investigate the global transcriptional landscape of soybean in different tissues and conditions. We identified biological processes and metabolic pathways that are enhanced or repressed in particular tissues and pointed TF families that probably control their expression, including candidates for downstream experiments (e.g. ectopic expression). We predicted the function of 93 unannotated hubs using a guilt-by-association approach. Further, we used this large GCN to gain insight into the fate of duplicated genes. We demonstrate that most soybean duplicated genes diverge in their expression profiles. However, duplicate pairs originating from WGDs are more often preserved in the same module than those arising by other mechanisms, suggesting an increased conservation of gene function for WGD-derived copies. We also demonstrate that duplicate pairs in the same module are likely under stronger purifying selection, as revealed by their lower Ka/Ks ratios. Collectively, our findings reveal potentially novel regulators and enlighten our understanding of the fate of duplicated genes during the evolution of the soybean genome.

## MATERIAL AND METHODS

### Data collection and processing

The normalized expression data (TPM, Transcripts Per Million) for 1,298 samples were downloaded from the recently published Soybean Expression Atlas [16]. Samples were filtered by using the standardized connectivity (Z.k) method in the R package *WGCNA* [27]. For that, we constructed a sample network based on squared Euclidean distances and assessed the overall scaled connectivity (Z.k) of the network. This method is efficient in filtering outlying samples and has been largely used in gene co-expression analyses [28-30]. Samples with Z.k < -2.5 were considered outliers and were filtered out. The resulting 1,284 samples represent 15 different tissues (Supplementary Figure S1). Further, to reduce noise, we excluded genes with TPM < 5 in more than 20% of the samples. Finally, the 30,563 × 1,284 matrix was log_2_ transformed by using log (TPM+1).

### Network reconstruction and module detection

The signed network was inferred by using the R package *WGCNA* [27]. We assessed the scale-free topology fit to choose the most suitable β power. The lowest β power for which the network reached a scale-free topology (R^2^ = 0.8) was 17 (Supplementary Figure S2). Pairwise correlations between genes were calculated by using the Pearson Correlation Coefficient (PCC). The PCC matrix was then transformed to an adjacency matrix, which was used to construct a topological overlap matrix (TOM). The TOM considers both PCC and shared neighbors to determine similarity, which amplifies disparity between gene pairs, making it effective for module detection. Genes were clustered into modules with hierarchical clustering based on dissimilarity (1 – TOM). Finally, we calculated pairwise similarity between modules and merged the ones that had PCC ≥ 0.8. Minimum module size was set to 30. The R package *igraph* (http://igraph.org) [31] was used for extraction and visualization of subgraphs.

### Hub gene inference and functional enrichment

Hub genes are those with the highest number of connections (i. e. degree) in a network. Intramodular connectivity (kWithin) was used as degree to detect intramodular hubs. The choice of thresholds for hub identification is somewhat arbitrary. While some groups consider the top 10% genes with the highest degree as hubs [32, 33], others use module membership (MM) to define such genes [34]. MM is a measure of how close a given gene is to its module, and it can be calculated by correlating a gene to the module eigengene. Here, we combined both parameters and considered hubs the top 10% genes with the highest intramodular connectivity and with MM ≥ 0.8. Enrichment analysis for GO terms, pathways and conserved protein domains was performed in PhytoMine (https://phytozome.jgi.doe.gov/phytomine/begin.do), and REVIGO was used to remove redundancy [35]. Soybean TFs were downloaded from PlantTFDB [36], and TF enrichment was determined by performing an one-sided Fisher’s exact test with the R package *bc3net* [37]

### Analysis of upstream cis-regulatory elements

We searched for regulatory motifs in the regions between -1000bp and +200bp relative to the transcription start site, as done in *A. thaliana* [38]. These regions were extracted with *bedtools* [39]. A *de novo* motif discovery was performed by using MEME [40]. As background, we used an order 0 Hidden Markov Model (HMM) with all expressed genes, using the script *fasta-get-markov* available in MEME suite.

### Simulated networks, paralogous genes and selection analysis

We simulated 10,000 degree-preserving networks in R through successive permutations of node labels to verify the probability of finding our results by chance. Paralogous gene pairs derived from dispersed, tandem, proximal, transposed, and whole-genome duplications were download from the Plant Duplicate Gene Database [41]. We used Ks thresholds to subdivide WGD-derived pairs into the WGDs that happened 13 mya (Ks ≤ 0.4) and 58 mya (Ks > 0.4). For simplicity, a smaller number of pairs potentially arising from older WGD events were combined in this latter category. Ka, Ks and Ka/Ks ratios were computed using the Perl script *calculate_Ka_Ks_pipe*.*pl*, available in [41]. Paralog pairs in the same module were interpreted as having similar transcriptional profiles. Pairs were interpreted as having different transcriptional profiles if: i. genes were distributed in different modules; ii. only one gene from the pair was present in the network or; iii. both genes were in the *grey* module, which stores genes that were not assigned to modules.

## RESULTS

### Degree centrality analysis reveals that hubs are enriched in essential genes

After hierarchically clustering the genes based on TOM dissimilarity and merging the modules with 80% or more similarity, we generated a GCN containing 29 modules (Figure 1; Supplementary Table S1). The largest and smallest modules were *chocolate3* (n = 6,250) and *steelblue3* (n = 30), respectively. The *grey* module, which stores unassigned genes, had 4,766 genes. We selected the top hub from each module and analyzed their functions in PhytoMine. Top hub genes tend to encode proteins of paramount importance, such as succinate dehydrogenase (Glyma.19G08600), an RNA polymerase subunit (Glyma.06G217400), DNA topoisomerase (Glyma.14G137100), and cellulose synthase (Glyma.06G324300) (Supplementary Table S2).

**Figure 1.**
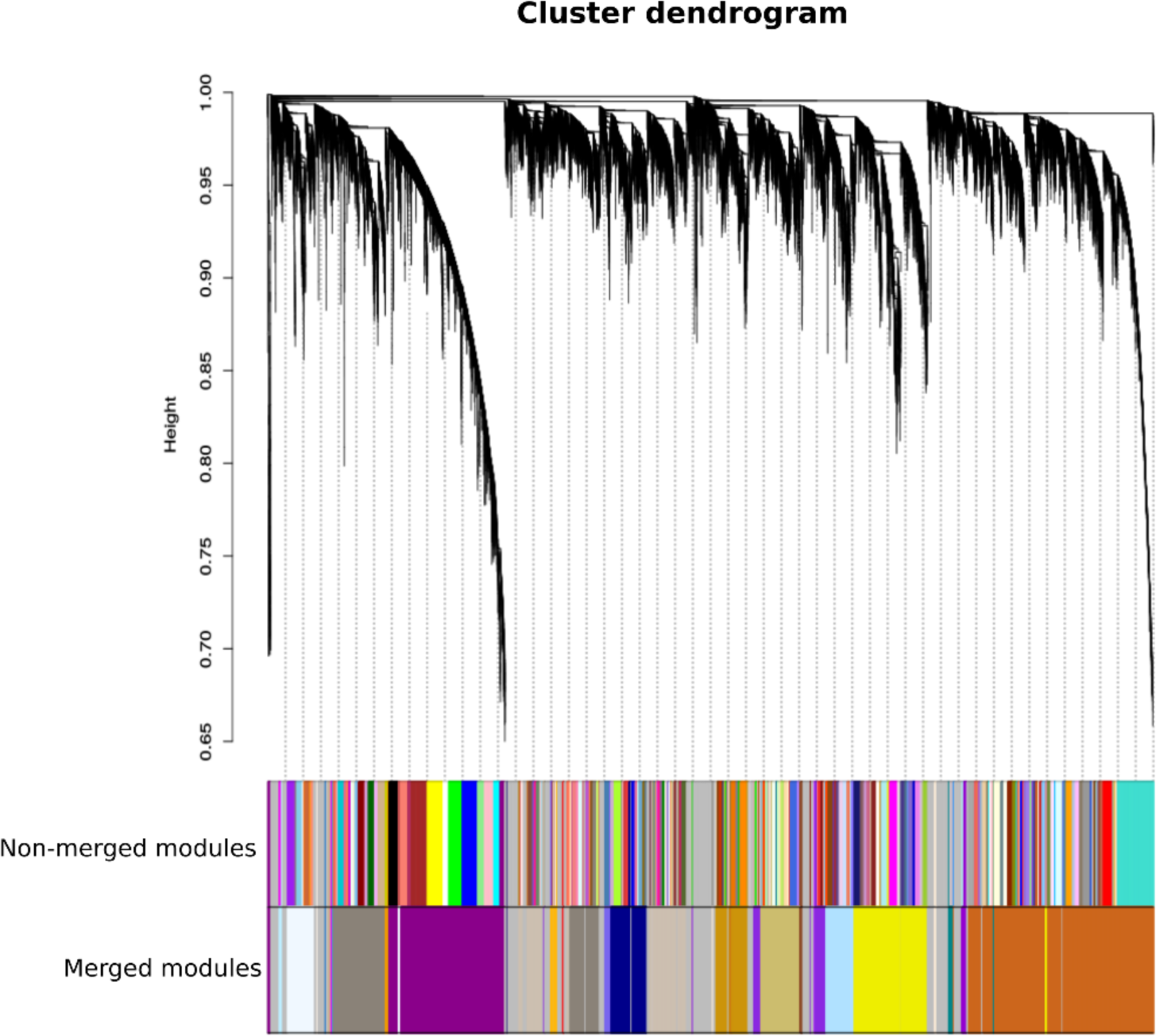
Dendrogram and module colors before and after merging similar modules. In the WGCNA standard pipeline, each module is given a color. Modules were merged based on similarity of their eigengenes. The module eigengene is the first principal component of a principal component analysis, which summarizes the expression of the module. Modules with eigengenes more than 80% similar (r = 0.8) were merged.

Jeong *et al* found that hubs tend to be essential in yeast (*S. cerevisiae*) protein-protein interaction networks [42]. We hypothesized that degree centrality in a GCN is also associated with gene essentiality in soybean. Essential genes in plants are often referred to as embryonic lethal genes (i. e. genes that compromise embryo viability when not expressed). *A. thaliana* essential genes have been diligently catalogued over the years by Meinke’s group, who recently published a curated list of 510 *EMBRYO-DEFECTIVE (EMB)* genes [43], out of which 481 have soybean orthologs (1,010 soybean genes). Importantly, the soybean orthologs of the essential genes (Supplementary Table S3) are significantly overrepresented among GCN hubs (Fisher’s exact test; p = 5.8e-05). Further, essential genes that are hubs are enriched in functions related to nucleic acids metabolism, translation and regulation of cell division, such as replication factors, ribosomal proteins, and elongation factors.

Next, we investigated if any of our modules was individually enriched for essential genes and essential hub genes. The modules *darkmagenta, yellow2, lightskyblue1, lightblue1, chocolate3, darkgoldenrod3*, and *darkviolet* were significantly enriched in essential genes (Fisher’s exact test; p = 7.8e-05, 2.3e-19, 4.5e-02, 5.5e-09, 7.8e-06, 5.8e-09, and 6.0e-03, respectively). A functional enrichment analysis revealed that, respectively, these modules are related to photosynthesis and gluconeogenesis; translation and ribosome biogenesis; RNA synthesis and processing; RNA metabolism; DNA replication; protein ubiquitination and regulation of transcription and; *darkviolet* had the enriched protein domain acetyl-CoA carboxylase alpha subunit. The latter three modules were also enriched with essential hubs (Fisher’s exact test; p = 9.6e-08, 1.3e-04, 4.8e-02, respectively). These findings show that these gene clusters are essential to embryo viability and could be experimentally validated in the future. Overall, our findings corroborate the well-established role of hub genes as essential players of the cell physiology [44, 45].

### Module hubs uncover biological processes associated with specific tissues

We selected the hub genes from each module and performed an enrichment analysis for GO terms, pathways and protein domains (see methods for details). Information on the number of hubs per module are in Supplementary Table S4. Only 14 of 29 modules (48.3%) had enriched GO terms, pathways or protein domains, and we identified the function of nine of the 14 modules based on enriched biological processes (GO-BP) and pathways (Supplementary Table S5). Further, we identified module-tissue relationships by correlating each module eigengene to each of the 15 tissues available in our dataset (Figure 2; Supplementary Table S6), some of which are discussed below.

**Figure 2.**
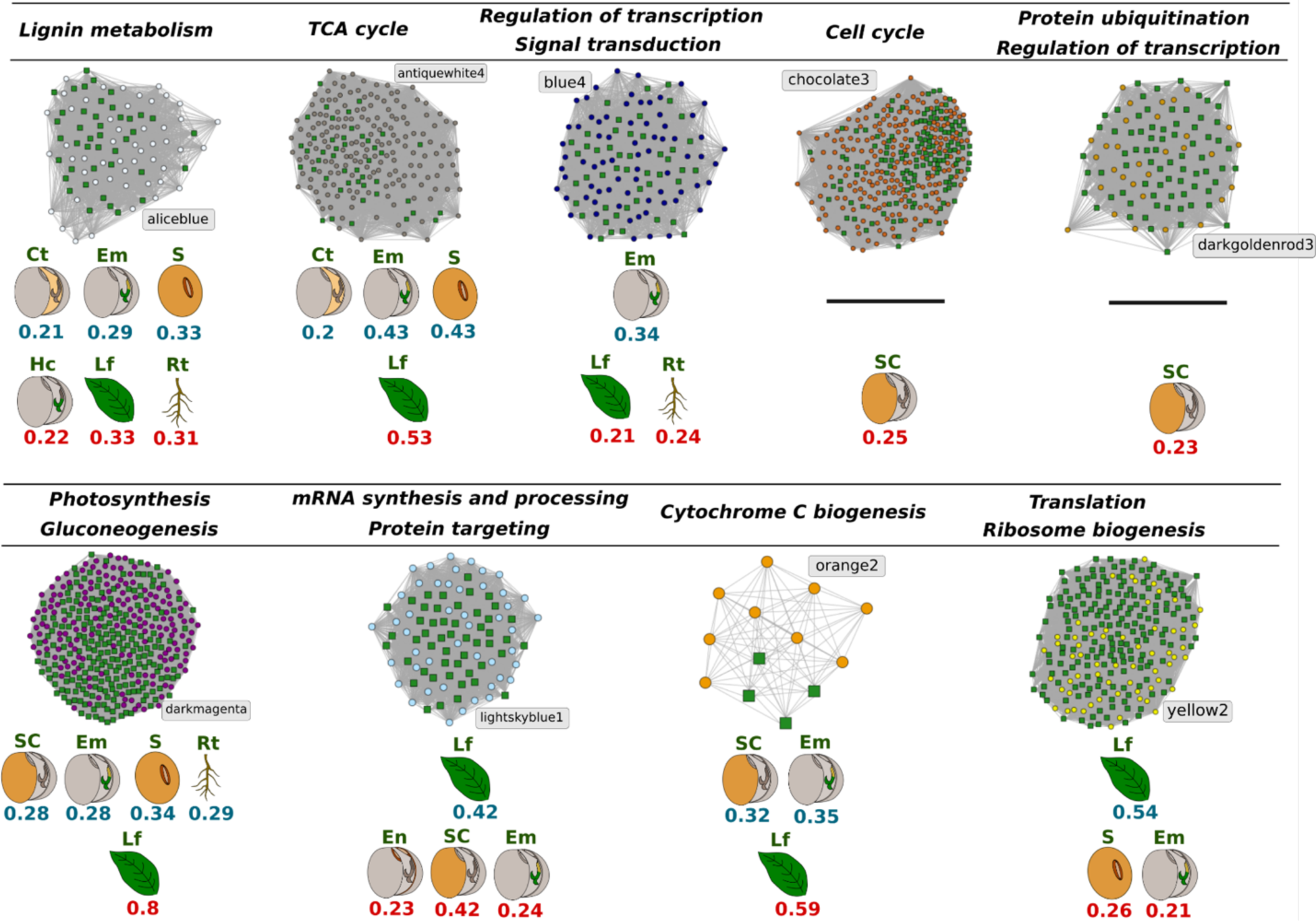
Hubs from modules with enriched biological processes (GO) and pathways. For each subgraph, green squares represent hubs that are TFs. Below each subgraph, we illustrate tissues where the genes tend to be down-regulated (blue) and up-regulated (red). Up-and down-regulation in each tissue were obtained by calculating Pearson correlation coefficients between module eigengenes and a binary matrix *m*_*ij*_ containing 1 if the sample *i* corresponds to the tissue *j*, and 0 otherwise. P-values were derived by using the function corPvalueStudent() in WGCNA. Given the heterogeneity of the data, correlations were considered significant if r ≥ 0.2 and p < 0.05. Ct, cotyledon; Em, embryo; S, seed; Hc, hypocotyl; Lf, leaf; Rt, root; SC, seed coat; En, endosperm.

Genes in the module *aliceblue* are related to lignin and phenylpropanoid metabolism, such as laccases (Glyma.01G112600, Glyma.03G077900, Glyma.08G359100), peroxidases (Glyma.01G163100, Glyma.11G049600, Glyma.11G078400, Glyma.12G195500), xylem cysteine proteinase (Glyma.04G014700, Glyma.06G014800), and cytochrome b561 (Glyma.02G306800). Module-tissue correlation showed that they are up-regulated in leaf, root, and hypocotyl (r = 0.33, 0.31, 0.22; p = 1e-34, 3e-29, 1e-15, respectively) and down-regulated in cotyledons, embryo and seeds (r = -0.21, -0.29, -0.33; p = 1e-14, 8e-26, 2e-34, respectively). Lignins are polymers made up of monolignols, which are polymerized into lignin by laccases and peroxidases [46]. Because of their strong covalent bonds to cellulose in secondary cell walls, lignins provide mechanical support to plant organs such as stems and leaves, which explains the up-regulation of *aliceblue* genes in this tissue [47]. In addition, lignins are difficult for herbivores to digest, constituting a defense mechanism. Lignin is also well known for its role in the stability of vascular tissues such as xylem [48]. The xylem cysteine proteinase encoded by Glyma.04G014700, also in *aliceblue*, is likely involved in the autolysis of tracheary elements during xylogenesis in roots [49]. Lignin is primarily synthesized in hypocotyls during seedling establishment, as shown in growing hemp hypocotyls [50]. Since lignin biosynthesis begins in the hypocotyl at the seedling stage, such genes are also expected to be down-regulated in embryo, seeds and cotyledons, which are mostly represented by developing seeds in our dataset. Thus, our results recapitulate some of the physiological and anatomical roles of lignin.

Hub genes in the module *antiquewhite4* encode proteins related to the tricarboxylic acid (TCA) cycle and aerobic respiration, such as isocitrate dehydrogenase (Glyma.13G144900, Glyma.10G058100, Glyma.16G118900) and ureide permeases (Glyma.01G058500 and Glyma.02G116300). These genes are up-regulated in leaf (r = 0.53; p = 5e-93) and down-regulated in seed, embryo and cotyledon (r = -0.43, -0.43, -0.2; p = 7e-60, 6e-60, 2e-13, respectively). As most seed-related tissues in our dataset represent developing seeds, these genes are expected to be down-regulated, since seed reserves are accumulated during development. Storage proteins and other nutrients are then mobilized during germination to promote early seedling development [51]. The up-regulation of TCA cycle-related genes in leaf is likely because the TCA cycle is one of the major sources of redox equivalents for oxidative phosphorylation, which provides ATP for sucrose synthesis [52].

In *blue4*, we found genes related to transcriptional regulation and signaling, such as ABC transporters (Glyma.13G361900, Glyma.07G015800), ferrochelatase (Glyma.04G050400), phosphatases (Glyma.06G290200, Glyma.08G071900), kinases (Glyma.04G088800, Glyma.06G090800, Glyma.13G073900, Glyma.14G001300, Glyma.02G311200), TFs (Glyma.03G127600, Glyma.02G232600, Glyma.07G023300, Glyma.14G200200, Glyma.09G005700, Glyma.12G212300, Glyma.13G289400), and adenylate kinase (Glyma.10G048200). *Blue4* genes were slightly up-regulated in leaf and root (r = 0.21 and 0.24; p = 6e-14, 5e-18, respectively) and down-regulated in embryo (r = -0.34; p = 6e-36). We analyzed the annotation of *blue4* orthologs in *A. thaliana* and found that this module is likely related to defense against pathogens, especially fungi. Among these orthologs, we found the ABC transporters PEN3 and PIS1 (AT1G59870 and AT3G53480), which participate in salicylic acid-dependent resistance against direct pathogen penetration and in secretion of phenolic compounds involved in Fe uptake [53]. Absorbed Fe ions can then be used by the ferrochelatase FC1 (AT5G26030) to supply heme for defensive hemoproteins [54]. Similarly, the TFs WRKY33 and TGA6 from this module are involved in the activation of salicylic acid-responsive genes and pathways mediating response to fungal pathogens, respectively [54, 55]. Innate immune responses rely on the detection of microbe-associated molecular patterns (MAMPs) by pattern recognition receptors (PRRs). Most *A. thaliana* PRRs are leucine-rich repeat (LRR)-receptor kinases [56, 57]. *Blue4* also harbor orthologs of *A. thaliana* innate immunity genes, such as an LRR-receptor kinase (AT5G01950) and an interleukin-1 receptor-associated kinase (AT1G14370). We also found an ortholog of the receptor kinase CERK1 (AT3G21630), which is involved in response to chitooligosaccharides, a very common plant MAMP [56, 58]. Hence, the up-regulation of these genes in leaf and root is coherent, whereas their down-regulation in embryo is likely because of its physical protection by other organs and tissues [59].

Hubs from *darkmagenta* encode proteins related to photosynthesis and gluconeogenesis, such as phosphoribulokinase (Glyma.01G010200), glycine transaminase (Glyma.01G026700), photosystem II enhancer protein 1 (Glyma.02G061100), NADPH-dependent thioredoxin reductase 3 (Glyma.02G185300), fructose-bisphosphate aldolase (Glyma.04G008300), glyceraldehyde-3-phosphate dehydrogenase (Glyma.04G015900), and fructose-1,6-biphosphatase (Glyma.07G142700). These genes are remarkably up-regulated in leaf (r = 0.8; p = 7e-292) and down-regulated in embryo, root, seed, and seed coat (r = -0.28, -0.29, -0.34, and -0.28; p = 4e-25, 2e-26, 4e-36, 2e-25, respectively). This is expected because in leaves, photosynthesis is responsible for CO_2_ fixation in triose phosphate, which feeds sucrose synthesis [60]. Hence, our findings support a strong transcriptional co-regulation of sucrose synthesis and photosynthesis genes. In root and seed-related parts, however, the down-regulation of these genes is due to their very low photosynthetic potential as compared with other plant parts, making them predominantly heterotrophic [61-64].

Finally, hubs from *orange2* are enriched in genes involved in cytochrome C biogenesis and assembly (e.g. Glyma.06G254100, Glyma.12G147300). This module is up-regulated in leaf (r = 0.59; p = 2e-119) and down-regulated in seed-related parts (embryo and seed coat, r = -0.35, -0.32; p = 1e-37, 6e-32, respectively). The *Arabidopsis* ortholog (AT5G54290) of these genes encodes CcdA, a thylakoid membrane cytochrome C biogenesis protein that transfers reducing equivalents from stromal thioredoxins to the thylakoid lumen [65]. Many proteins located in the thylakoid lumen depend on redox regulation, and it has been shown that the *Arabidopsis ccda* mutant is incapable of assembling the cytochrome b_6_f in thylakoids [66].

### Functional prediction of unannotated hubs

Guilt-by-association (GBA) approaches have been widely used for function inference [67-70]. In the context of GCN, it assumes that genes with similar expression patterns are co-regulated and more likely to share the same pathway or biological process. We explored the GCN and found 106 unannotated (i.e. genes without brief description in PhytoMine) hubs, of which 93 were distributed in the nine modules for which we could assign some biological function, as described above. By using a GBA approach, we predicted functions for all these 93 hubs (Supplementary Table S7), which might unveil interesting players in soybean transcriptional regulation.

We also found that 28 out of 29 modules had annotated top hubs. The top hub in the module *antiquewhite4* (Glyma.19G074200) encoded a protein of unknown function. We performed a functional enrichment analysis of its direct neighbors (r ≥ 0.7) and found that they are involved in response to auxin (p = 1.19e-3), which is likely the function of this unannotated gene. These genes encode auxin-responsive GH3 family proteins (Glyma.03G256200 and Glyma.19G258800), and SAUR-like auxin-responsive proteins (Glyma.04G231100, Glyma.06G134000, Glyma.08G004100, Glyma.19G254000).

The soybean genome remains far from being completely annotated. While 80.71% of soybean genes have some, although often minimal, functional annotation in PhytoMine’s brief description, the remaining 19.29% encode proteins of unknown function. Network-based approaches like the one reported here might help us understand the functions of many interesting genes, in particular those potentially associated with agronomic traits. As hub genes are typically major players of the cell physiology, our results can be relevant for the soybean genomics community.

### Enrichment of transcription factor families uncovers major regulators of important biological processes

We applied a Fisher’s exact test and found that 10 of the 29 modules are enriched in TFs (Supplementary Table S8), suggesting that the functions involved in these modules require a more complex transcriptional regulatory architecture. We also found 6 modules enriched in specific TF families (FDR ≤ 0.05; Table 1). We hypothesize that the overrepresented TF families in each module have key regulators of the biological processes enriched in the module (Table 1). We also performed a *de novo* motif discovery in promoter regions of genes in each module with enriched TF families (Supplementary Figure S3), although none of the identified motifs matched those from the same family in JASPAR and other Arabidopsis databases available in Tomtom (MEME suite) [71]. Because of their regulatory roles, TFs are often reported as important targets for developing transgenic lines with increased efficiency in several biological processes and metabolic pathways [72-74]. Hence, the genes reported in Table 1 could be interesting targets for plant transformation studies.

**Table 1.** Overrepresented TF families in soybean co-expression modules. One-sided Fisher’s exact test with Benjamini-Hochberg correction was used. TF families were considered significantly overrepresented if FDR < 0.05.

| Module name | TF family | FDR | Module function | Gene ID |
| --- | --- | --- | --- | --- |
| aliceblue | WOX | 8.83e-4 | Lignin metabolism | Glyma.04G040900,<br>Glyma.14G084600 |
| antiquewhite4 | TALE | 1.27e-3 | TCA cycle and aerobic respiration | Glyma.09G095700,<br>Glyma.10G081700,<br>Glyma.12G188800,<br>Glyma.13G312900,<br>Glyma.17G104800 |
| blue4 | WRKY | 2.98e-6 | Response to biotic stress | Glyma.02G232600,<br>Glyma.07G023300,<br>Glyma.08G218600,<br>Glyma.09G005700,<br>Glyma.12G212300,<br>Glyma.13G289400,<br>Glyma.14G200200,<br>Glyma.15G003300,<br>Glyma.15G110300 |
| darkmagenta | CO-like | 7.28e-3 | Photosynthesis and gluconeogenesis | Glyma.05G233700,<br>Glyma.07G091400 |
| lightskyblue1 | MYB-related | 5.41e-4 | mRNA synthesis and processing | Glyma.09G257700,<br>Glyma.13G128500,<br>Glyma.15G236400,<br>Glyma.18G234900 |
| yellow2 | C2H2 | 1.04e-2 | Translation and ribosome biogenesis | Glyma.11G189500,<br>Glyma.12G084700,<br>Glyma.12G181400 |

### Network topology and the possible fate of soybean duplicated genes

We sought to explore the constructed GCN to explore the fates of duplicate genes. All soybean duplicated gene pairs were downloaded from Plant Duplicate Gene Database [41]. Duplicated gene pairs were classified in tandem, proximal, transposed, dispersed and whole-genome duplication (TD, PD, TRD, DD and WGD, respectively). Based on the Ks distribution, WGD-derived duplicates were subdivided into WGD 58 mya and WGD 13 mya, representing WGD events that happened 58 and 13 million years ago, respectively (Supplementary Figure S4). The co-occurrence of duplicates in modules can help understand their fates after duplication, as each module has a unique expression profile (Supplementary Figure S5). In 27.2% of the gene pairs (22,003/80,946), both genes were excluded during data filtering (see methods for details) and are thus not present in the network, while in 27.6% (22,355/80,946) only one gene of the pair was present in the network. Finally, both genes were present in the network for 45.2% of the pairs (36,589/80,946). Expectedly, these three categories have distinct gene expression levels (Supplementary Figure S6).

We analyzed the co-occurrence of duplicated genes and the prevalence of their modes of duplication in modules. Duplicate pairs in which both genes were absent in the network were not considered. The co-occurrence of duplicate pairs in the same module was interpreted as an indicative of functional conservation, as they reflect similar expression profiles after duplication. On the other hand, when duplicate pairs were in different modules or when only one gene from the pair was present, we interpreted it as an indicative of functional divergence. In order to assess the significance of these results, we created 10,000 degree-preserving simulated networks (Figure 3; see methods for details). We observed that most pairs displayed divergent expression profiles or signs of fractionation (Figure 3). Strikingly, WGD pairs, particularly those from the most recent WGD, showed greater levels of retention when compared to other modes of gene duplication, both in the same and in different modules. These observations are possibly correlated with previous reports on the increased retention of WGD duplicates involved in intricate systems (e.g. regulatory genes) and protein complexes [23].

**Figure 3.**
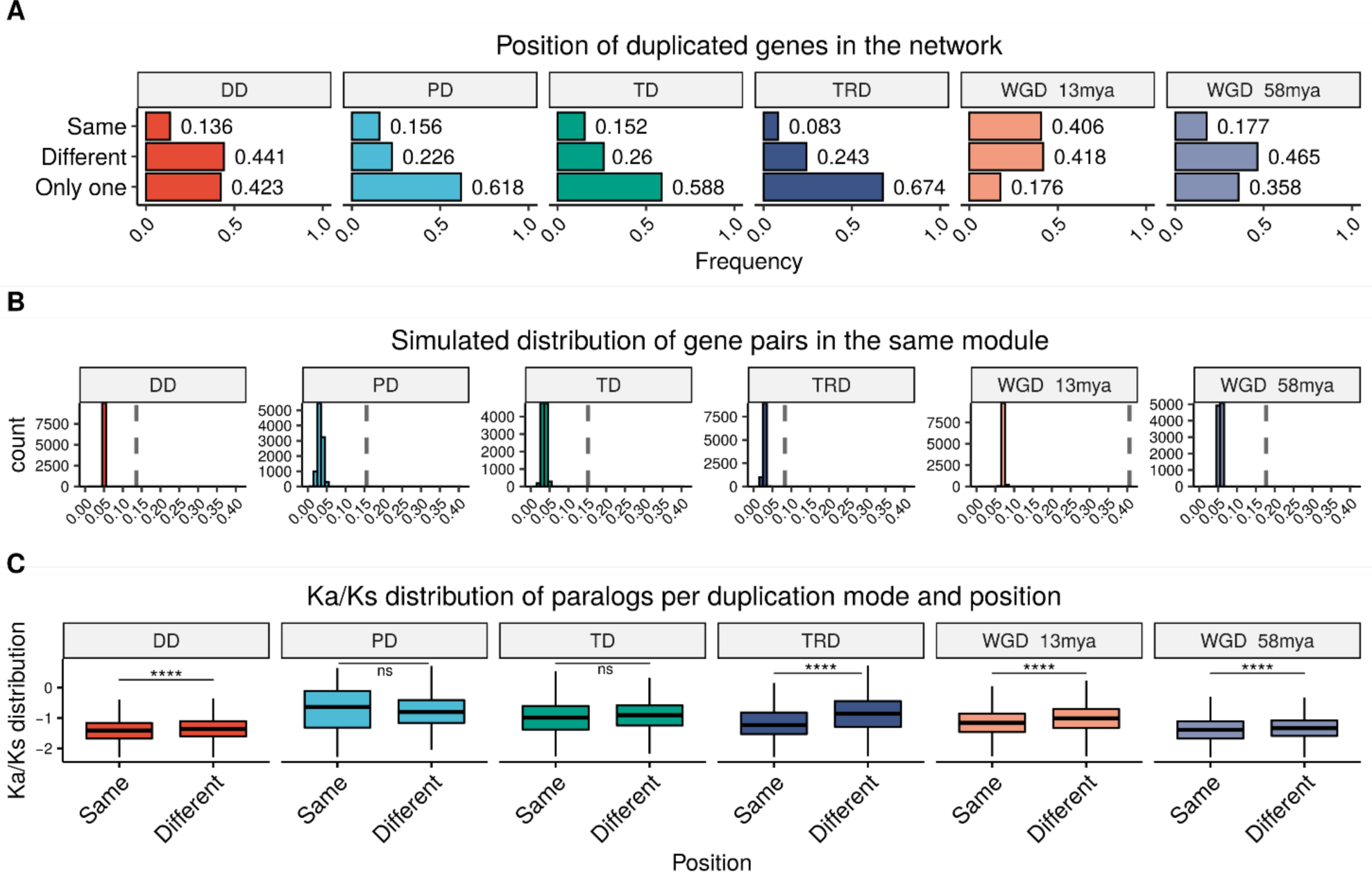
Distribution and Ka/Ks analysis of duplicate pairs in the network. A. We observed that the minority of the duplicate pairs derived from all modes of duplication were in the same module in the GCN, supporting functional divergence as the most common fate of duplicated genes. B. We created 10,000 degree-preserving simulated networks by permuting the node labels of the GCN. Next, we compared the distribution of frequencies in which duplicate pairs were in the same module (histograms) with that observed in the real GCN (dashed lines). When both genes from a pair fell into the *grey* module, we treated them as being in different modules, as *grey* stores unassigned genes. C. Distribution of Ka/Ks ratios per mode of duplication and presence in the same module. Ka/Ks values were log transformed using log(Ka/Ks + 0.1) for better visualization. Except for proximal and tandem duplication-derived gene pairs, Ka/Ks ratios are significantly lower for paralog pairs within the same module, supporting that they are under stronger purifying selection at both transcriptional and sequence levels. DD, dispersed duplications; PD, proximal duplications; TD, tandem duplications; TRD, transposed duplications; WGD, whole-genome duplications; ns, no significant difference; ****, p<0.0001.

As previously shown, subfunctionalization is by far the most common fate of retained duplicated genes [75], although it is not totally clear how this process happens at the transcriptional level. Our findings suggest that the most common fate of duplicated genes is functional divergence, as 59,4% to 91.7% of the duplicates do not belong to the same module. Nevertheless, the frequency of genes in the same module is higher than the expected by chance, supporting that the transcriptional similarity of part of the duplicates is under selective pressure. This finding provides new insights into how retained copies are regulated throughout the soybean genome.

Finally, we hypothesized that paralog pairs with similar expression profiles (i.e same module) are under stronger purifying selection as compared to paralog pairs with different expression profiles (i.e. members of the pair in different modules, both in *grey* module or only one gene from the pair present in the network). We observed that, except for proximal and tandem duplication-derived pairs, paralog pairs with similar expression profiles have significantly lower Ka/Ks ratios as compared to those with different expression profiles (Wilcoxon test; p < 0.05; Figure 3C). This result suggests that duplicates that are retained in the same module are under stronger negative selection, both at transcriptional and sequence levels. These pairs could be involved in systems that rely on tightly regulated stoichiometric balance of its components.

## CONCLUSION

Here, we employed a thorough GCN analysis to explore the systems-wide transcriptional landscape of soybean tissues. We explored the functions of several modules and hubs using enrichment and GBA methods. Several of the identified modules are strongly associated with specific tissues, giving important clues about their physiological roles. We also found modules enriched in TFs, which probably correspond to systems dependent on more intricate transcriptional regulation architectures. Finally, we used our GCN to explore evolution of duplicate genes in soybean and found that most of the retained duplicated genes tend to diverge in expression, although genes from WGD are more often found in the same modules than those duplicated by other mechanisms. The GCN reported here provides insights into the roles of several soybean genes and co-expression modules, which might be important not only in the elucidation of gene functions, but also in exploring genes involved in relevant agronomic traits.

## AUTHOR CONTRIBUTIONS

Conceived the study: FA-S, KCM and TMV. Data analysis: FA-S, KCM and FBM. Funding, project coordination and infrastructure: TMV. Manuscript writing: FA-S and TMV.

## ACKNOWLEDGEMENTS

This work was supported by funding from Coordenação de Aperfeiçoamento de Pessoal de Nível Superior (CAPES, Finance code 001), Conselho Nacional de Desenvolvimento Científico e Tecnológico (CNPq), and Fundação Carlos Chagas Filho de Amparo à Pesquisa do Estado do Rio de Janeiro (FAPERJ). The authors declare no conflict of interest.

