## Supplementary figures for "Exploring the complexity of soybean (*Glycine max*) transcriptional regulation using global gene co-expression networks"

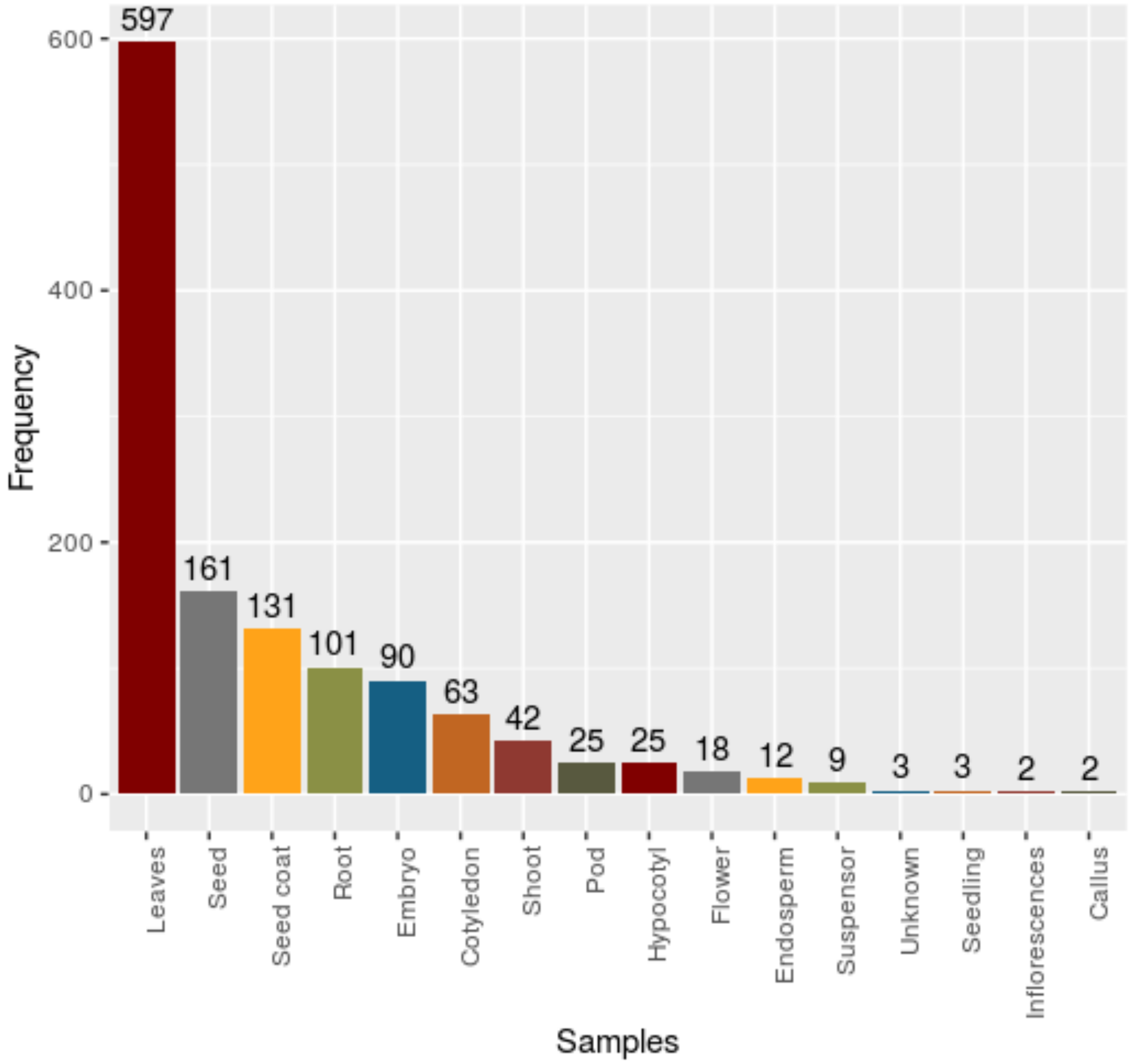


**Supplementary Figure S1. Absolute frequency of samples per tissue.** Fourteen samples downloaded from the Soybean Expression Atlas were considered outliers (Z.k < -2.5) and were then excluded from our analysis prior to network inference.

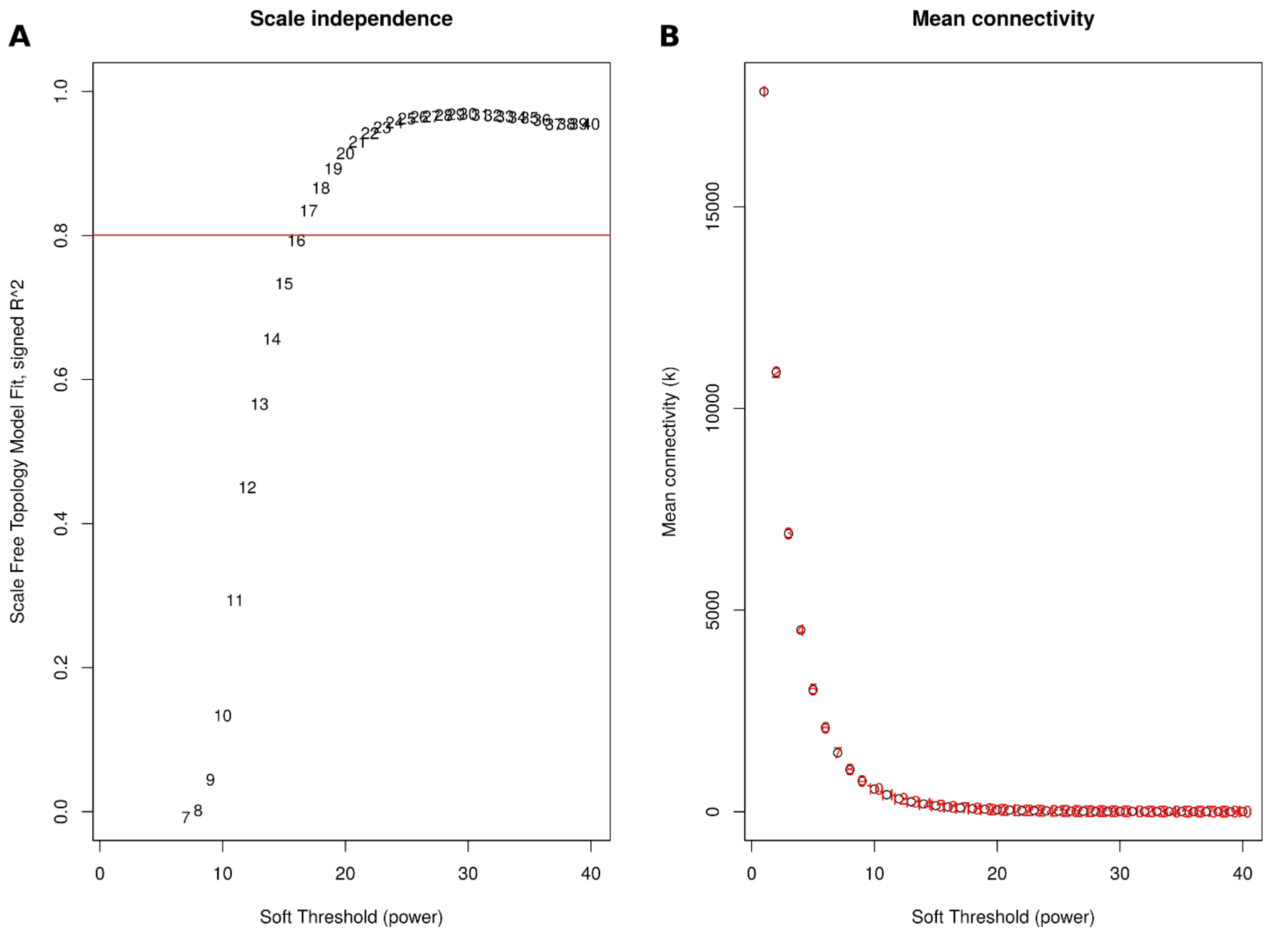


**Supplementary Figure S2. Scale-free topology fit.** As shown in A, a higher β implies a higher proximity of the network to a scale-free topology. On the other hand, a higher β leads to a lower number of connections (*k*) per node in the network (mean connectivity), as shown in B. Thus, choosing a β power is a trade-off between scale-free topology fit and mean connectivity. The lowest β power for which the network resembles a scale-free topology ($R^2$=0.8) and has a considerable mean connectivity was 17.

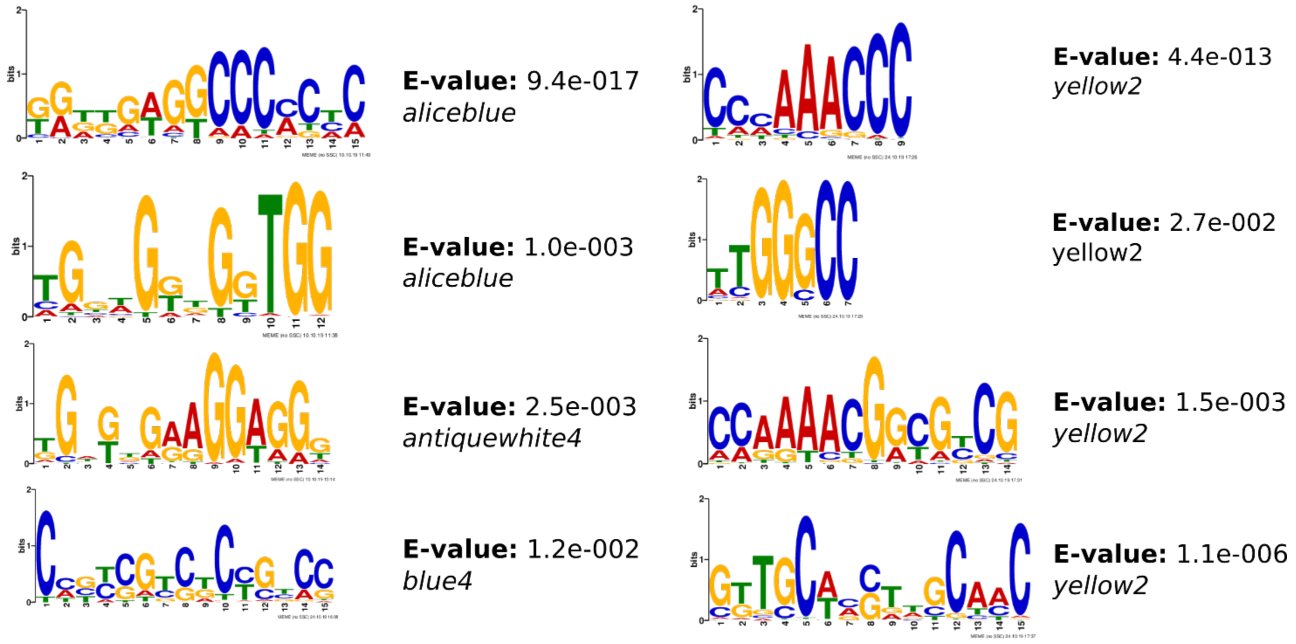


**Supplementary Figure S3. Motifs identified in promoter regions of genes in modules enriched for specific TF families.** Low-complexity regions were masked. A 0$^{th}$ order Hidden Markov Model of all expressed genes was used as background. The required motif site distribution was “zoops” (zero or one site per sequence). Minimum and maximum motif widths were set to 5 and 15, respectively.

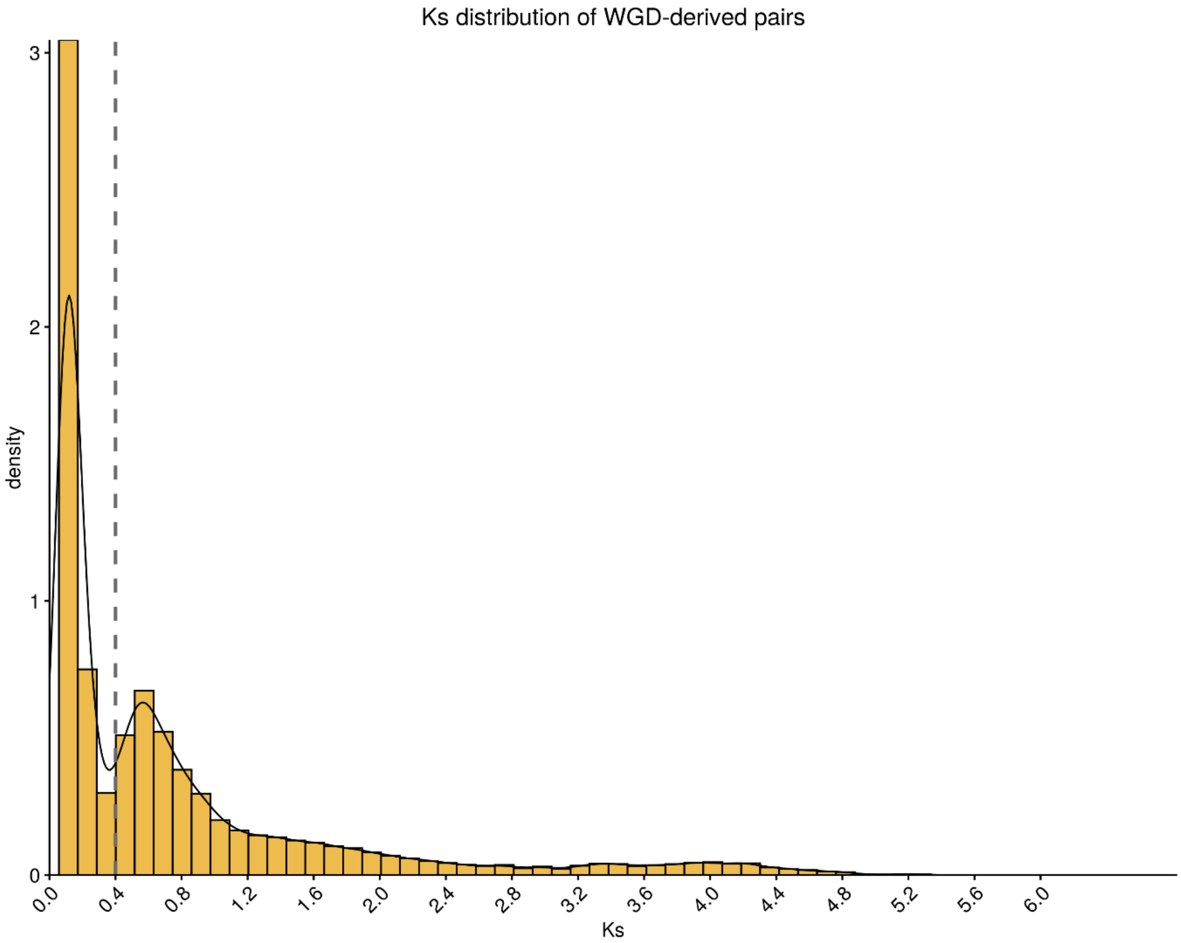


**Supplementary Figure S4. Ks distribution of WGD-derived duplicate pairs.** The two distinct peaks represent different whole-genome duplication events. WGD pairs with Ks rates up to 0.4 were classified as duplicate pairs that arose in the most recent soybean WGD (13 million years ago), while pairs with Ks rates higher than 0.4 arose in the most ancient soybean WGD (58 million years ago).

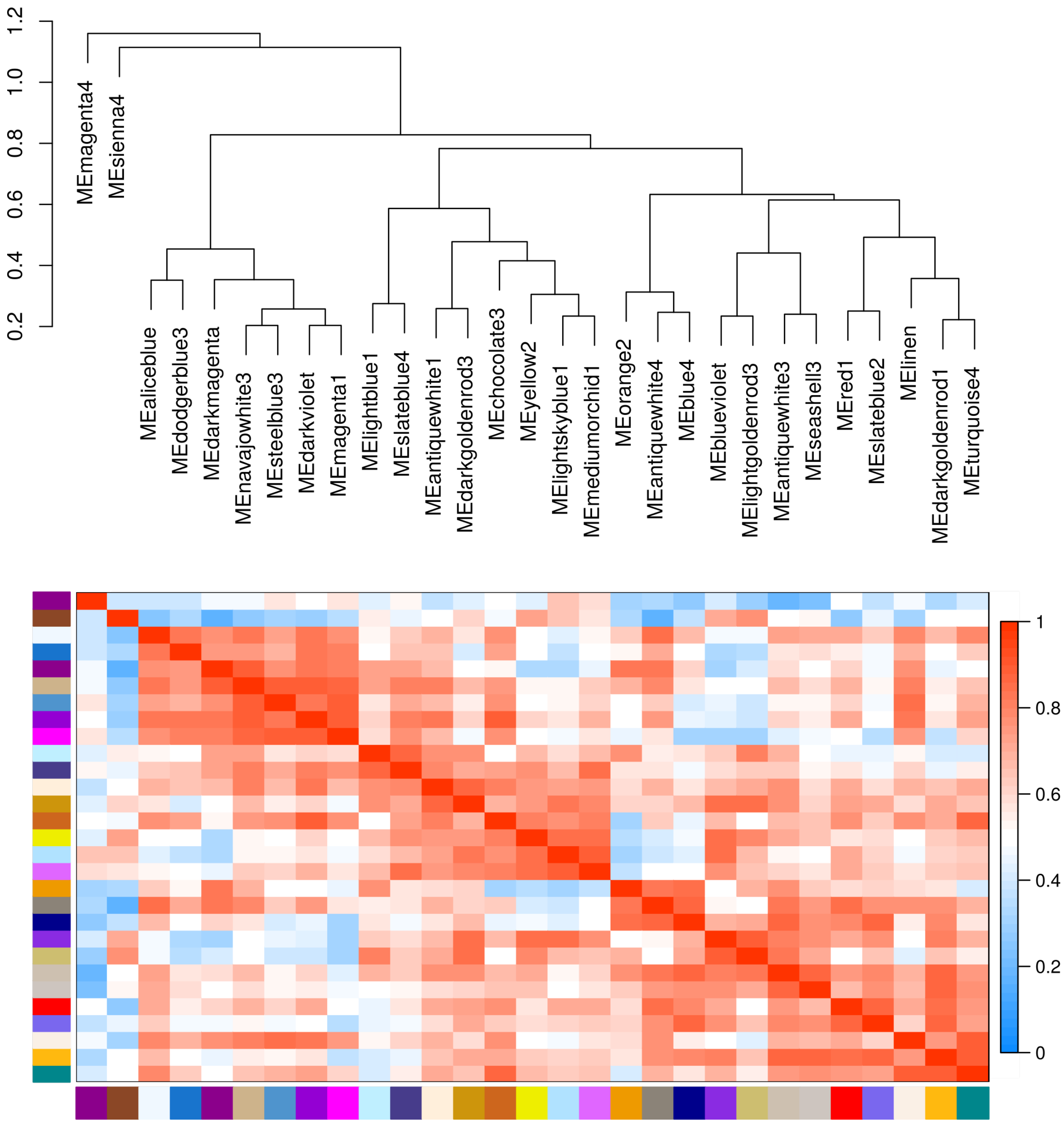


**Supplementary Figure S5. Eigengene networks.** The dendrogram was created by using the hclust() function in R with 1 – correlation of module eigengenes.

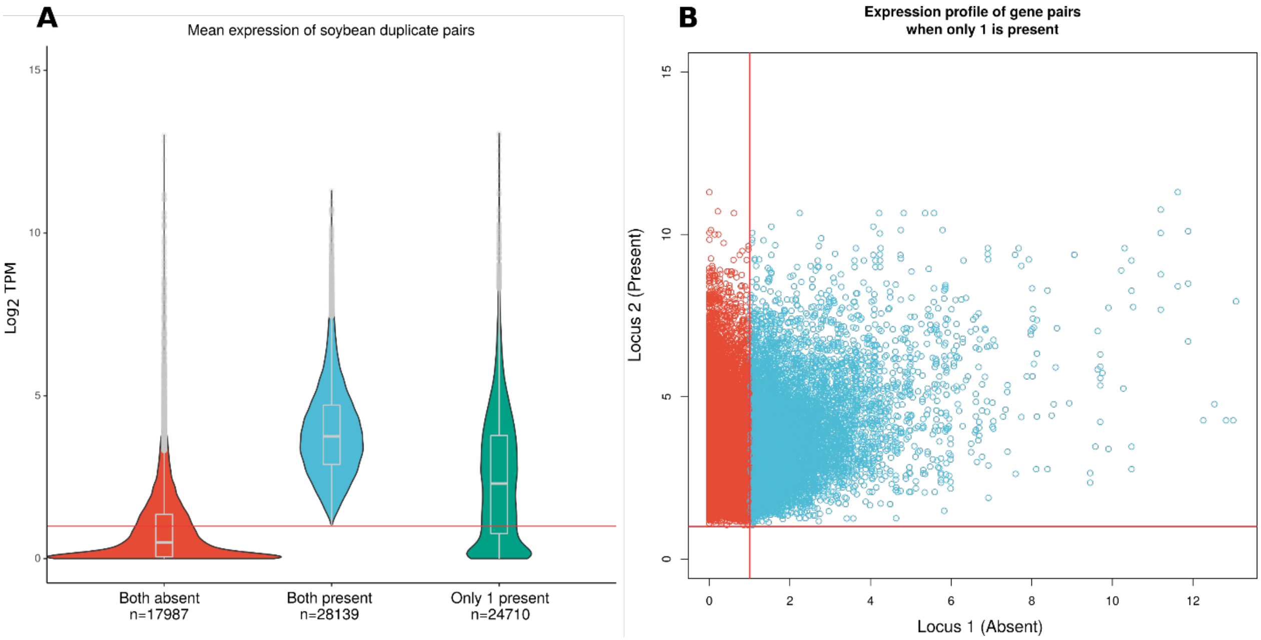


**Supplementary Figure S6. Expression patterns of soybean paralogous genes**. A. Mean expression of genes from duplicate pairs that were absent in the network, duplicate pairs that are included in the network and duplicate pairs for which only one gene was present. Gene pairs for which both genes were absent in the network were not used in our analysis. Most of the duplicate pairs that were excluded during pre-processing have mean expression lower than 1. All duplicate pairs in the network have mean expression greater than 1 and follow a Gaussian distribution. Duplicate pairs for which only one gene was present range from no expression to high expression values. However, the genes with high expression values are probably highly expressed in a small number of samples, as the mean is considerably affected by outliers. B. Mean expression of duplicate pairs for which only one gene is present in the network. All genes from *locus* 2 (present) have mean expression greater than 1. In *locus* 1 (absent), however, only 50.4% of the genes have mean expression greater than 1. As previously pointed, the ones that had mean expression much greater than 1 are probably affected by outliers.
